# A human iPSC-derived motor neuron-myogenic cell coculture platform to evaluate neuromuscular junction innervation after axon injury and in Spinal Muscular Atrophy

**DOI:** 10.1101/2025.10.17.682712

**Authors:** Keunjung Heo, Xiangsunze Zeng, Kathleen Zhang, Kuchuan Chen, Shannon Zhen, Anika Naveen, Richard M. Giadone, Rasheen Powell, Roshan Pandey, Lee L. Rubin, Clifford J. Woolf

**Author notes:** Correspondence (Clifford J. Woolf). These authors contributed equally (Keunjung Heo, Xiangsunze Zeng).

## Abstract

Traumatic nerve injury is challenging as motor neurons with damaged axons repair slowly, which can lead to muscle degeneration, while in Spinal Muscular Atrophy (SMA), muscle innervation is reduced. While pro-regenerative or neuroprotective compounds have been identified, their specific ability to enhance or restore neuromuscular junction (NMJ) function in patients remains unclear due to a lack of in vitro human models that track axon growth, NMJ innervation and muscle function. We developed a human iPSC-derived motor neuron-myogenic coculture platform that enables real-time monitoring of axon growth NMJ innervation and axon regeneration and muscle activity following axonal injury, and for SMA-derived motor neurons. We identified spontaneous synchronized GCaMP6f muscle activity as a useful functional marker of NMJ formation. Using this platform, we show that blebbistatin, a pro-regenerative non-muscle myosin II (NMII) inhibitor differentially regulates growth cone dynamics in injured versus uninjured motor neurons, resulting in enhanced NMJ reinnervation. This highlights the therapeutic potential of developing pro-regenerative compounds to promote NMJ innervation. We also confirm that NMJ function is reduced in SMA type 0 (prenatal onset) and type I (pediatric onset) patient-derived motor neurons in the coculture. This human stem cell-based framework can be used, therefore, to evaluate pro-regenerative compounds for axon injury and neuroprotective one for neurodevelopmental and neurodegenerative disorders.

## INTRODUCTION

Traumatic nerve injuries are relatively common, but once motor axons are damaged, effective full regeneration is challenging. Following peripheral nerve injury, extracellular calcium rises, activating Schwann cells, immune cells, and fibroblasts, which migrate to the lesion and form a bulb-like seal (1-4). Once sealed, axons retract, calcium levels drop, and actomyosin-driven cytoskeletal remodeling as well as transcriptional changes initiate regeneration, while a loss of neuronal input causes rapid myofiber atrophy (5-8). Although previous studies have identified pro-regenerative compounds that promote neuronal regrowth, their ability to restore neuromuscular junction (NMJ) formation in humans remains largely unknown, as no in vitro human models currently exist to assess both regeneration and NMJ function over time after axonal injury (9-14), or in motor neuron diseases.

Although the mechanisms driving axon regeneration and NMJ formation are fundamentally distinct, compounds have rarely been tested in complex multicellular systems. As a result, evaluating potential efficacy largely depends on single cell cultures or in vivo animal models. These challenges are even greater in the context of neurodegenerative diseases, where uncertainties about disease onset complicate therapeutic testing. To overcome these challenges, it is critical to move beyond single-cell culture approaches and develop in vitro human cell based multicellular coculture platforms with an unbiased analysis. Such systems could enable a more accurate evaluation of pro-regenerative or neuroprotective compounds and offer a greater potential for identifying therapeutic strategies for restoring motor neuron and muscle function.

To deal with these challenges, we developed a human iPSC-derived motor neuron-myogenic coculture platform that enables high-throughput, real-time monitoring of NMJ innervation and muscle function as well as axon regeneration. By combining a spatially defined culture system with laser-induced axotomy, GCaMP6f-based calcium imaging, and automated analysis, we can quantify spontaneous synchronized muscle activity as a functional readout of NMJ formation. Using this system, we tested the effects of the non-muscle myosin II (NMII) inhibitor blebbistatin, a pro-regenerative compound (15-18), on NMJ function. A previous study showed that NMII inhibition at the site of nerve injury enhances regeneration and motor recovery, but its systemic use is limited by off-target adverse effects (17, 19-21). Three days of blebbistatin treatment after an axotomy in the coculture system increased the number of activated myogenic cells. Mechanistically, we found that NMII acted differently in naïve versus axon injury conditions which must be due to its distinct roles at growth cones.

We further applied this platform to examine changes in NMJ function for spinal muscular atrophy motor neurons following the initial motor neuron innervation of target muscles. Although SMA is traditionally classified as a neurodegenerative disorder, emerging evidence suggests it also involves neurodevelopmental defects (22-26). Using our coculture system, we found that spontaneous synchronized muscle activity was reduced in myogenic cells cocultured with motor neurons derived from SMA type I patients, a pediatric-onset form of the disease, despite no obvious defects in neuronal outgrowth during early NMJ innervation. In contrast, SMA type 0, antenatal-onset form, neurons exhibited reduced neurite growth over time, likely contributing to the decreased numbers of activated myogenic cells and synchronized events. These findings indicate impaired NMJ formation and activity in SMA during early development and identify a critical time window for testing effective neuro-enhancing compounds. This coculture platform enables, then, investigation of the efficacy of pro-regenerative and neuroprotective compounds on NMJ function in both axonal injury and neurodegenerative disease contexts and revealed new insights into the role of NMII on NMJ innervation.

## RESULTS

### NMJ-driven spontaneous synchronized muscle activity captured in a motor neuron-myogenic co-culture platform

To establish a motor neuron (MN)-myogenic cell co-culture system (Fig. 1A), we utilized Isl1- and ChAT-positive MNs (Fig. 1B) and myogenic cells that express structural markers muscle myosin, and multinucleated sarcomeres (Fig. 1C) Coculture was initiated by seeding MNs onto MHCK7-GCaMP6f-expressing myogenic cells (Fig. 1A). Following innervation and NMJ formation, postsynaptic sites in the myogenic cells were identified by α-bungarotoxin (BTX) staining (Fig. 1D). At this stage, two distinct spontaneous GCaMP6f activity patterns were observed: synchronized activity, a population-based muscle activity characterized by simultaneous activation of multiple myogenic cells, and sporadic activity resulting from asynchronous calcium transients in individual cells (Fig. 1E). To quantify the number of synchronized myogenic cells, we used CaImAn an unbiased, automated approach for calcium imaging (27) in 1 min recordings acquired at 10 Hz, measuring both the number of activated cells per frame and the number of synchronized events. In myogenic cell-only cultures, GCaMP6f activity was sporadic with no synchronization (Fig. 1E-1G). However, when cocultured with MNs, the myogenic cells exhibited synchronized GCaMP6f activity, and this was abolished following treatment with the acetylcholine receptor (AChR) blocker curare (Fig. 1E-1G). Synchronization was defined as a single event in which more than eight myogenic cells were activated simultaneously per frame (Fig. 1F) which occurred on average at 1.9 times during the 1 min imaging period (Fig. 1G). A temporal analysis showed that synchronized activity emerged by day 2 following MN innervation and persisted through day 4 of co-culture, and then gradually declined thereafter (Fig. 1H, and 1I). The frequency of this coordinated myogenic activation peaked on day 4 of coculture (Fig. 1J), suggesting that the rise in the number of activated cells serves as an early indicator of NMJ functionality before an increase in synchronized events is observed.

**Figure 1.**
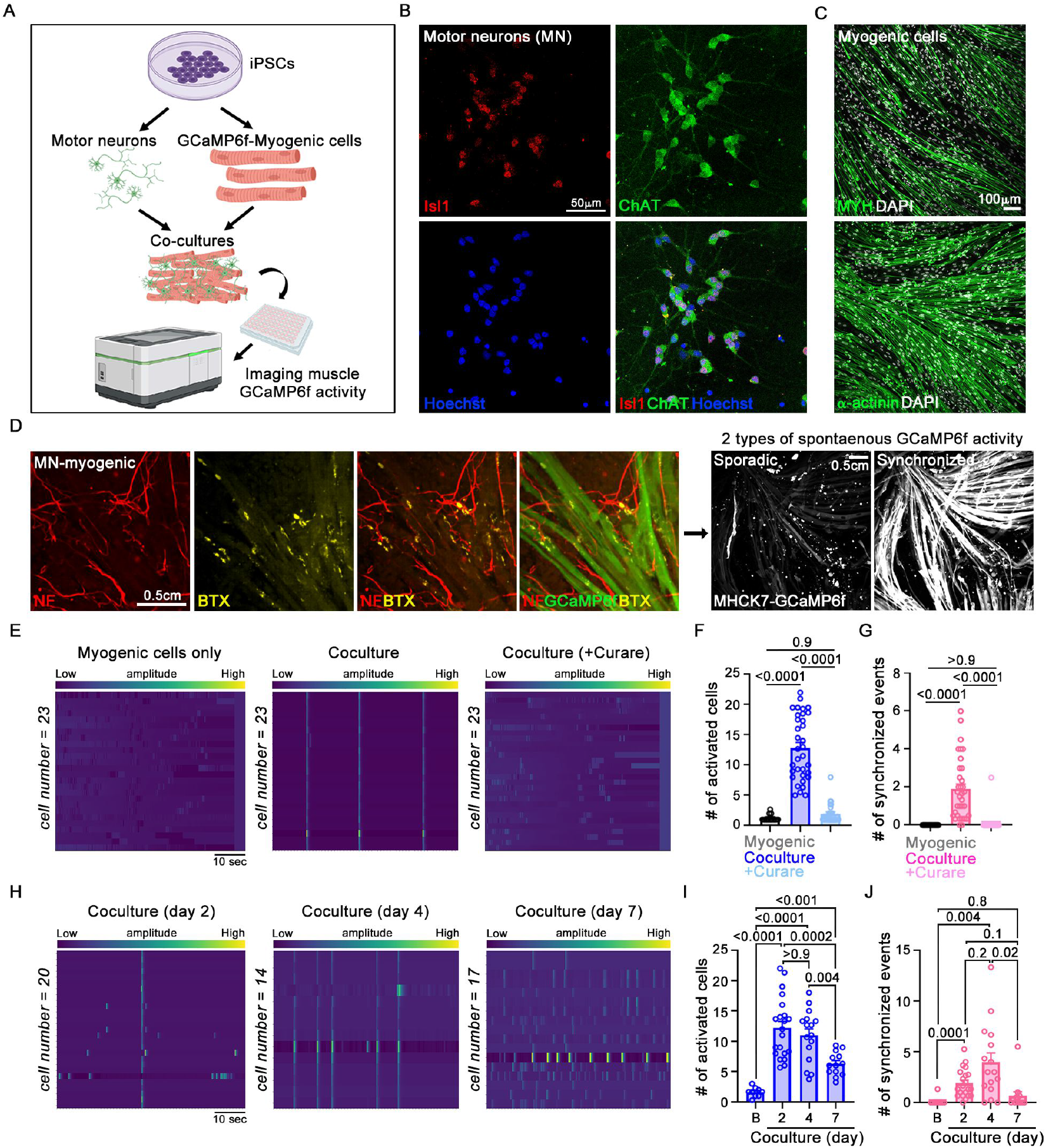
Motor neuron-skeletal myogenic cell coculture system to assess NMJ formation and function. Experimental schematic: iPSCs from a single donor were differentiated into MNs and MHCK7-GCaMP6f-expressing skeletal myogenic cells, co-cultured, and muscle activity recorded following innervation, using a high-throughput imaging platform. **(B)** MNs stained with motor neuron markers, Isl1, and ChAT with nucleic acid dye Hoechst. Scale bar, 50 μm. **(C)** Skeletal myogenic cells labeled with fast-twitch muscle myosin marker, MYH1 or sarcomere marker, α-actinin, and DAPI. Scale bar, 100 μm. **(D)** MN-myogenic cocultures labeled with neurofilament (NF), GCaMP6, and α-bungarotoxin (BTX), showing sporadic and synchronized GCaMP6f activity. Scale bar, 0.5 cm. **(E)** Heatmaps of GCaMP6f activity (1 min at 10 Hz) in myogenic-only, MN-myogenic, and 50 μM curare-treated cocultures. **(F and G)** Quantification of synchronized activity; number of activated myogenic cells per frame **(F)**, and number of synchronized events per min **(G)** across myogenic-only (n=12), MN-myogenic coculture (n=34); and curare-treated coculture (n=27) conditions. Data represents three biological replicates. Stats: One-way ANOVA with Tukey’s post hoc test, error bars: SEM. **(H-I)** GCaMP6f analysis of MN-myogenic cocultures over time. **(H)** Heatmaps of muscle GCaMP6f activity recorded in MN-myogenic cocultures at day 2, 4, and 7. Quantification of synchronized muscle activity: number of active myogenic cells exhibiting synchronization **(I)**, and number of events per min **(J)** in baseline (n=9), day 2 (n=21), day 4 (n=16) and day 7 (n=13) cocultures. Data represents at least three biological replicates. Stats: Welch’s ANOVA with Dunnett’s T3 post hoc test, error bar: SEM.

### Spontaneous synchronized muscle activity re-emerges following axon injury in motor neuron-myogenic cell cocultures

To assess NMJ functionality after axon injury using synchronized activity as a readout of NMJ reinnervation, we established a co-culture platform by seeding MN spot cultures within myogenic cells, enabling a spatial separation between the two cell types for a laser-cut axonal injury (Fig. 2A). Synchronized GCaMP6 activity was monitored over time after a laser-cut axonal injury in the same region used for pre-injury NMJ innervation assessment (Fig. 2B). This approach enabled direct comparison of the same GCaMP6f-expressing cells before and after injury. We confirmed that the GCaMP6 activity is at the same location where axons form NMJs with myogenic cells, by labeling the axons with neurofilament and the postsynaptic sites with BTX staining to label AChR clustering in the myogenic cells (Fig. 2C). Before injury, synchronized activity was detected, typically involving five myogenic cells (Fig. 2D), with an average frequency of 0.5 events per minute (Fig. 2E). The synchronized activity with the MN spot cultures (Fig. 2D and 2E) involved fewer myogenic cells than the 2D coculture, likely due to the more localized distribution of the axons. The synchronized activity disappeared immediately following axon injury but began to re-emerge by day 3 post-injury with an increasing number of activated cells, resembling the pre-injury pattern, and remained robust through day 5 post-injury (Fig. 2D and 2E). By day 7 post-injury, this activity gradually declined (Fig. 2D and 2E), consistent with the pattern seen in uninjured cocultures.

**Figure 2.**
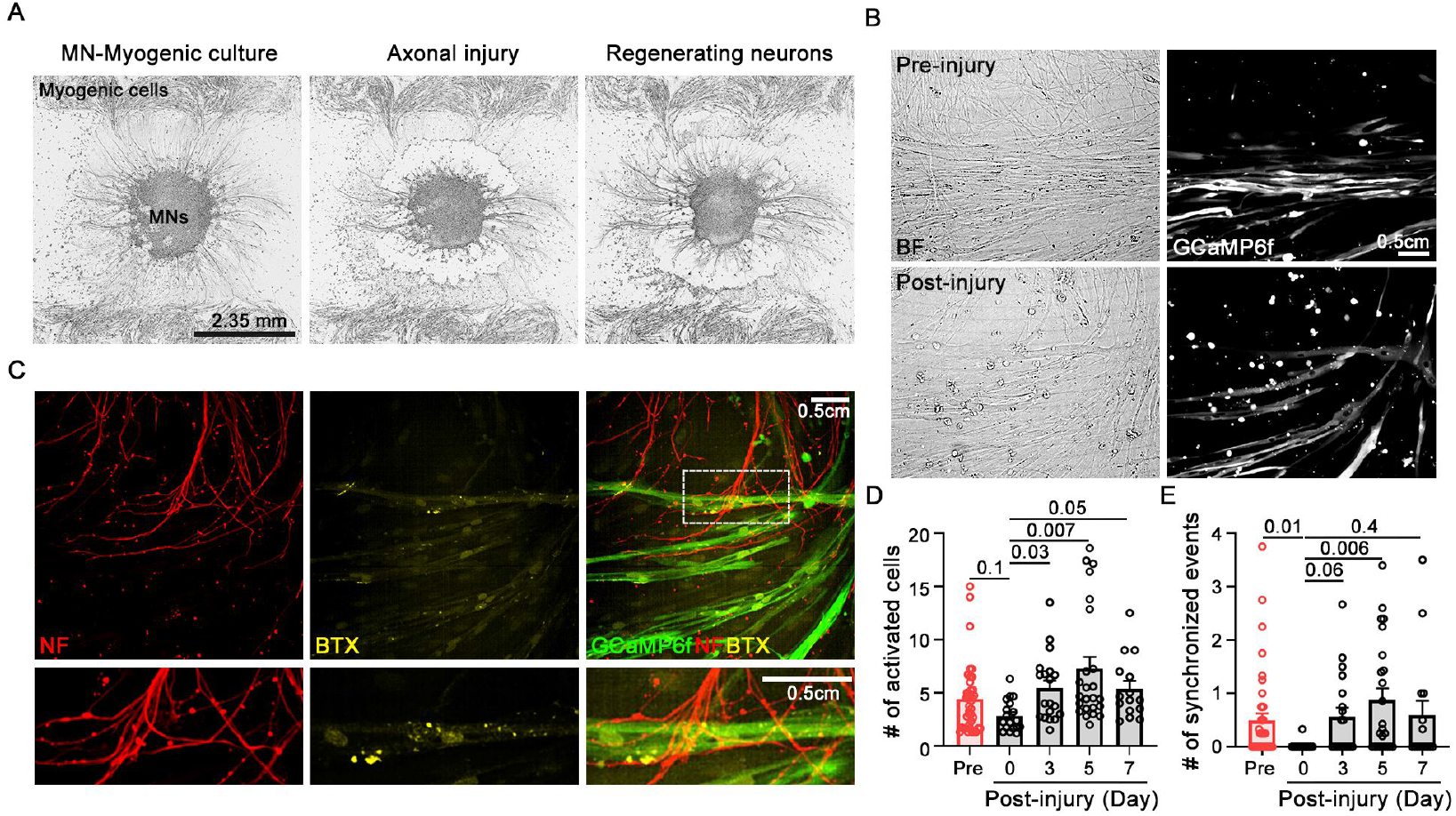
Synchronized GCaMP6f activity in muscle reflects functional synaptic innervation. **(A)** Skeletal myogenic cells co-cultured with spatially localized MN spot cultures. Following axon innervation of the myogenic cells, a laser-induced axonal injury was produced, and neuronal regeneration monitored. Scale bar, 2.35 mm. **(B)** Images of the cocultures before and after injury. **(C)** Neurons labeled with NF, muscle cells with GCaMP6f, and postsynaptic site with α-BTX. Scale bar, 0.5 cm. **(D and E)** Quantification of synchronized muscle activity: number of active myogenic cells **(D)** and synchronized events **(E)** pre-injury (n=40) and day 0 (n=18), day 3 (n=19), day 5 (n=24), and day 7 (n=15) post-axonal injury. Data represents three biological replicates. Stats: One-way ANOVA with Tukey’s post hoc test, error bars: SEM. *p*-values: indicated in each graph.

### Blebbistatin enhances NMJ connectivity after nerve injury

We evaluated the effects of the non-muscle myosin II (NMII) inhibitor blebbistatin on NMJ functionality after axon injury using this platform, given that it enhances recovery of muscle function after a nerve injury in vivo (17). Blebbistatin enhanced axon regeneration in the coculture system at day 2 post-axonal injury (Fig. 3A). To assess its impact on NMJ functionality, we examined the cultures on day 3 post-injury and showed an earlier complete restoration of NMJ function (Fig 3B), than after an axon injury without blebbistatin, which peaks on day 5 (Fig. 2G). Blebbistatin elevated the post- to pre-injury ratio of activated myogenic cells (Fig. 3B-3D), indicating that the enhanced axonal growth resulted in the quicker formation of functional NMJs. No significant changes in the number of synchronized events were observed, however, at this time. Consistent with these functional improvements, 3 days of blebbistatin treatment also enhanced the innervation of the myogenic cells compared to controls (Fig. 3E).

**Figure 3.**
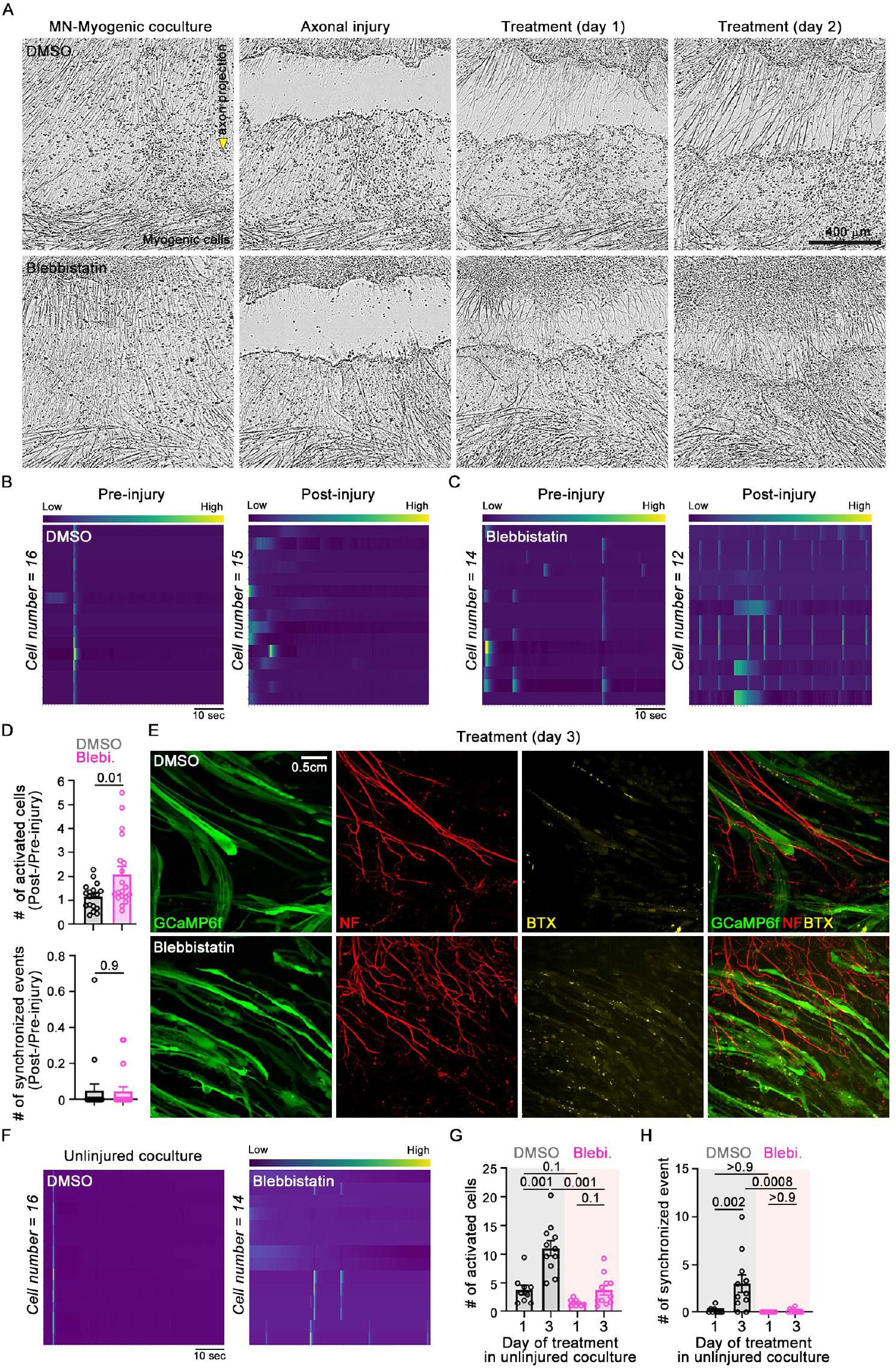
Blebbistatin enhances muscle GCaMP6 activity following axonal injury. **(A)** Images of MN-myogenic cocultures before injury, immediately after axonal injury, and at day 1 and day 3 following treatment with DMSO or 25 μM blebbistatin. Scale bar, 400 μm. **(B and C)** Heatmaps of muscle activity before injury and day 3 post-injury in DMSO **(B)** and blebbistatin **(C)** treated MN-myogenic cocultures. **(D)** Quantification of synchronized activity: number of active myogenic cells and synchronized events in DMSO (n=18), and blebbistatin (n=19) treated MN-myogenic cocultures. Data represents three biological replicates. Stats: Student’s *t*-test, error bars: SEM. *p*-values: indicated in each graph. **(E)** GCaMP6f expressing myogenic cells in cocultures stained with α-BTX and NF following treatment with DMSO and blebbistatin. Scale bar, 0.5 cm. **(F)** Heatmap of muscle activity in uninjured cocultures treated with DMSO or 25 μM blebbistatin. Quantification of synchronized muscle activity: number of active myogenic cells (**G**), and synchronized events (**H**). Sample sizes: day 1 DMSO (n=9), day 3 DMSO (n=11), day 1 blebbistatin (n=9), and day 3 blebbistatin (n=11). Data represents three biological replicates. Stats: Welch’s ANOVA with Dunnett’s T3 post hoc test for **G**, One-way ANOVA with Tukey’s post hoc test for **H**. Error bar: SEM, *p*-values: indicated in each graph.

To determine whether this effect was specific to the regenerative context, we next examined uninjured cocultures after the blebbistatin treatment. Unlike the enhanced activity observed in axonal injury model (Fig. 3D), blebbistatin treatment in uninjured cocultures decreased both number of activated cells and synchronized events compared to controls (Fig. 3F-3H). This reduction was detectable from day 1 of coculture and became even more pronounced by day 3 (Fig. 3G and 3H). These findings suggest that while NMII inhibition under naïve (uninjured) conditions promotes axonal growth many of the additional sprouts are nonfunctional and disrupt NMJ formation.

Although blebbistatin is classified as a non-muscle myosin II inhibitor, previous studies have reported that it can target muscle myosin II (28, 29). To investigate this, we evaluated myogenic cell morphology in myogenic-only cultures following blebbistatin treatment. In the absence of MNs, myogenic cells showed no spontaneous synchronized muscle activity on day 1 post-treatment, and this remained unchanged on day 3 (Fig. S1A-S1C). However, compared to vehicle-treated controls, blebbistatin treatment increased the number of activated cells and led to more frequent synchronization events at day 3 than at day 1 post-treatment (Fig. S1B and S1C). We next compared these findings with cultures treated with the skeletal myosin II inhibitor MPH-220 (30). Unlike blebbistatin, 3-day MPH-220 treatment did not affect the number of activated cells or the frequency of synchronized events in myogenic-only cultures (Fig. S1D and S1E). We further tested the muscle relaxant dantrolene, which inhibits the ryanodine receptor 1 by blocking intracellular calcium release (30, 31), and confirmed that it did not affect synchronized muscle activity (Fig. S1D and S1E). These findings indicate that the effects of blebbistatin on myogenic cells are not due to an off-target inhibition of skeletal muscle myosin II but reflect an unexpected effect on skeletal muscle function through a mechanism independent of intracellular calcium release and that blebbistatin both enhances axon reinnervation of NMJs after axon injury and increases activity of myogenic cells.

We next investigated whether the off-target effect of blebbistatin on myogenic cells influences NMJ formation and functionality. To address this, we assessed muscle morphology by labeling myogenic cells with the fast-twitch muscle myosin marker MYH1. 3-days of blebbistatin treatment reduced muscle thickness by 21.6% in uninjured cocultures (Fig. S1G). We then evaluated whether these morphological changes impacted NMJ formation by quantifying BTX-labeled postsynaptic sites. BTX intensity per cell remained unchanged after 3-days of blebbistatin treatment (Fig. S1F and S1G), indicating that blebbistatin reduces muscle thickness without disrupting AChR clustering.

### Early NMJ connectivity is impaired in myogenic cells cocultured with motor neurons derived from SMA type 0 and type I patients

Using this coculture platform, we evaluated whether early NMJ connectivity defects could be detected in a disease context. Spinal muscular atrophy (SMA) is traditionally classified as a neurodegenerative disorder caused by SMN1 gene mutations, leading to MN degeneration, muscle weakness, and multi-system dysfunction (26). However, recent studies suggest that the most severe forms, SMA type 0 and 1, may also link with neurodevelopmental dysfunction, as SMN protein appears to play a role in early brain development and cognitive function (23, 24, 32). To assess spontaneous muscle activity following axon innervation from MNs from an SMA background, we generated MNs from iPSCs derived from SMA type 0 (antenatal onset) and type I (pediatric onset) patients and co-cultured them with myogenic cells from a healthy donor (Fig. 4A). By day 2, control cocultures containing healthy donor-derived MNs (LiPSC-GR1.1) displayed a consistent increase in both the number of activated cells, and synchronized events compared to myogenic-only cultures (Fig. 4B-4D). In contrast, all cocultures with SMA patient-derived MNs showed reduced synchronized events (Fig. 4B and 4D), indicating that early NMJ connectivity was impaired at day 2. These results suggest that SMN deficiency may not only result in neurodegeneration but also impact the neurodevelopmental processes related to NMJ innervation.

**Figure 4.**
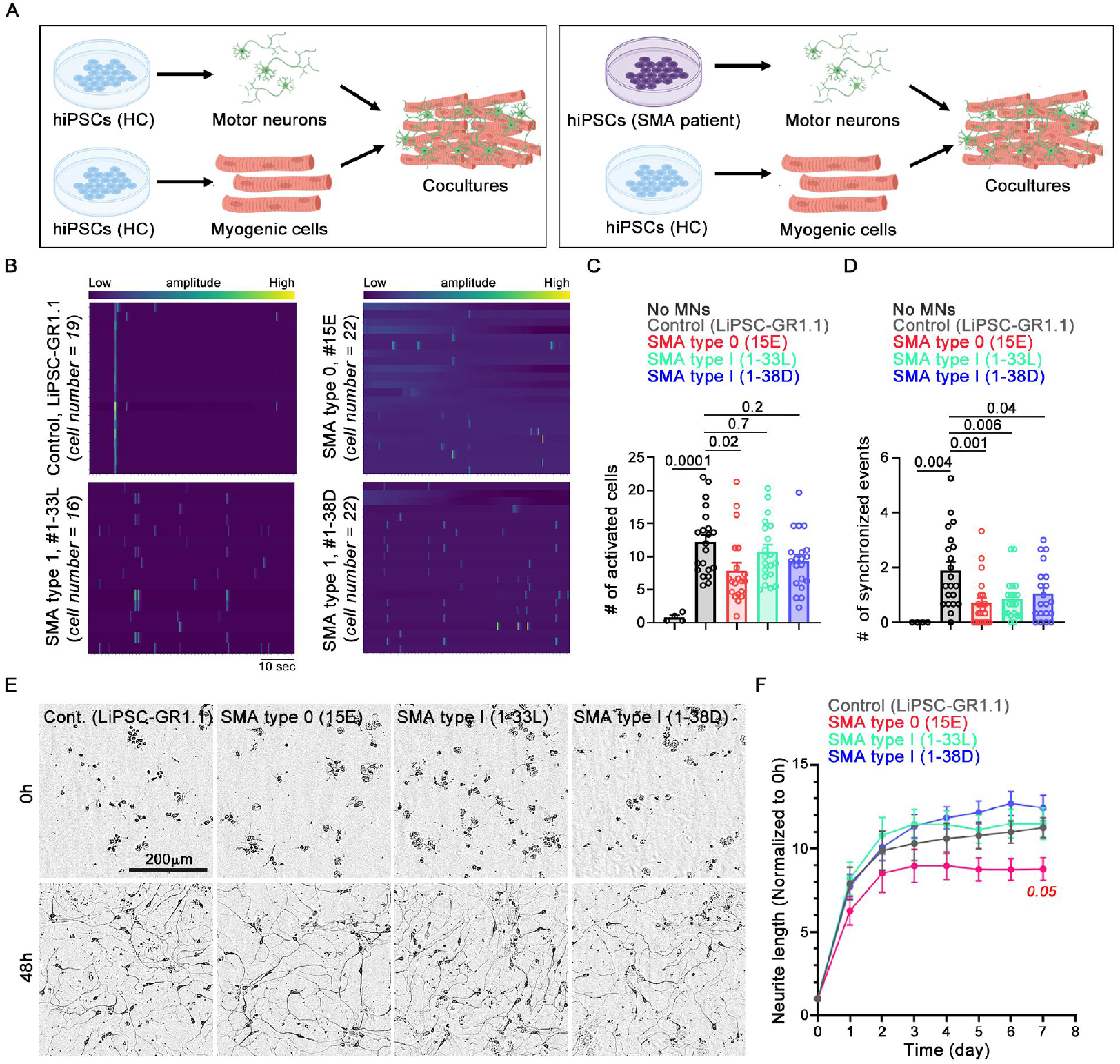
SMA type 0 motor neurons exhibit impaired NMJ connectivity. **(A)** Schematic illustrating the assessment of muscle GCaMP6f activity in an SMA background. MNs derived from healthy donor or SMA patient iPSCs were co-cultured with skeletal myogenic cells generated from healthy donor iPSCs. **(B)** Heatmap of GCaMP6f activity of myogenic cells cocultured with MNs from iPSCs from a healthy control on day 2 (LiPSC-GR1.1), SMA type 0 (15E), or SMA type I (1-33L and 1-38D) patients. **(C, D)** Quantification of muscle activity: number of active myogenic cells (**C**), and synchronized events (**D**) in day 2 cocultures. Sample sizes: myogenic cells with no MNs (n=4), with MNs from LiPSC-GR1.1 (n=21), 15E (n=20), 1-33L (n=20), and 1-38D (n=20). Data represents three independent biological replicates. Stat: One-way ANOVA with Tukey’s post hoc test, error bars: SEM, *p*-values are indicated in the graphs. **(E and F)** Neurite outgrowth in MNs derived from healthy control (LiPSC-GR1.1), SMA type 0 (15E), and SMA type I (1-33L, and 1-38D) iPSCs. **(E)** Representative images of MNs captured at 0, and 48h. Scale bar, 200 μm. **(F)** Time-course of neurite length, normalized to 0h time point. Data from at least three biological replicates. Sample sizes: LiPSC-GR1.1 (n=15), 15E (n=15), 1-33L (n=15), and 1-38D (n=14). Stats: Two-way ANOVA with Tukey’s post hoc test, error bars: SEM; Only significantly changed *p*-values are indicated (control vs. SMA type 0 at day 7: *p*=0.05).

Cocultures with SMA type 0 MNs consistently showed a decrease in the number of activated cells, whereas those with SMA type I MNs did not (Fig. 4B and 4C). We confirmed that the sporadic muscle activity in the SMA motor neuron coculture was blocked by curare (data not shown). To examine whether the decline in NMJ connectivity was linked to MN growth, we examined neurite outgrowth in MN-only cultures. MNs derived from an SMA type 0 patient showed a mild reduction in neurite outgrowth over time compared to those from healthy controls or SMA type I patients (Fig. 4E and 4F). In SMA type I, the reduction in NMJ activity appears to be independent of neuronal axon growth, in contrast, in SMA type 0 MNs which showed impairments in both NMJ activity and neuronal growth, resulting in a reduction in both the number of activated myogenic cells and synchronized events.

## DISCUSSION

This coculture platform enabled us to capture early NMJ innervation and activity. Spontaneous synchronized GCaMP6f muscle activity emerges once MNs innervate and form functional NMJs. The early activity peaked at day 2, with a more than 4-fold increase in the number of activated myogenic cells, followed by a peak in synchronized muscle activity at day 4. The activity then gradually declined, which may be a consequence of the absence of an interneuron input required to sustain spontaneous MN activity. Under physiological conditions, spinal interneurons coordinate MN firing and contribute to central pattern generator (CPG) activity, which is essential for rhythmic and coordinated motor output (27, 33-37). This platform nevertheless enabled an assessment of NMJ functionality using CaImAn, an AI-driven, open-source approach that automatically detects individual activated cells (27), which identified two key parameters: the number of activated myogenic cells per frame and the number of synchronized events per minute. Comparing neuronal regeneration after axon injury with and without blebbistatin exposure together, with observations on MNs from a SMA background, we found that the number of activated muscle fibers correlates with neurite outgrowth, likely reflecting NMJ formation, while the synchronized activity provides a measure of NMJ functionality.

Phenotypic pro-regenerative compound screens typically rely on neuron-only cultures to assess neurite growth. Our platform, where NMJ activity is the phenotype, enables us to evaluate the impact of pro-regenerative compounds on NMJ formation and function following a motor axon injury. We tested non-muscle myosin II inhibitor blebbistatin in the MN-myogenic coculture, providing a controlled system to evaluate drug effects on both motor neuron regeneration and NMJ functionality. We found that blebbistatin treatment increased the number of activated myogenic cells engaged in synchronized activity. This may reflect enhanced axonal branching (17), which could expand the pool of functional NMJs and activate more muscle cells. By contrast, in uninjured cocultures, blebbistatin reduced synchronized activity which indicates that while NMII inhibition under these conditions promotes axonal branching, many of these additional sprouts are likely nonfunctional. Thus, NMII inhibition appears to regulate axonal dynamics differently in regenerative versus naïve states.

The discrepancy between injured and naïve states raises two possible explanations. One is that NMII inhibition modulates growth cone dynamics differently during axon regeneration then in uninjured axon growth conditions. NMII localizes to actin rings in both axons and growth cones, where it modulates actomyosin force and microtubule dynamics (38, 39). After an axotomy, NMII inhibition disrupts the actin ring in growth cones, allowing microtubule extension (40-42), resulting in an increase in axonal density. However, in naïve uninjured conditions, NMII’s role in modulating actomyosin forces appears to interfere with NMJ formation and negatively influences muscle morphology. We also observed morphological alterations in myogenic cells after blebbistatin treatment that were independent of neuronal innervation, as these effects occurred in myogenic-only cultures. These alterations may result from blebbistatin’s effects on calcium signaling, which can influence early myogenic differentiation and maturation (5, 7, 8, 43-45). However, these effects did not disrupt AChR clustering, which is essential for NMJ activity, indicating that short-term blebbistatin treatment (3 days) in MN-myogenic cell cocultures does not impair NMJ formation. These findings highlight the importance of developing blebbistatin analogs with no skeletal muscle activity.

Using the MN-myogenic cell co-culture platform, we also showed that SMA type 0 shows a neurodevelopmental like defect, despite being conventionally regarded as a neurodegenerative disorder (23, 24, 32). We observed reduced synchronized activity in myogenic cells co-cultured with MNs derived from SMA type 0 or type 1 patients. A possible interpretation for this is that while initial NMJ formation occurs in the SMA background, they fail to fully mature, leading to diminished spontaneous synchronization events. These results underscore the significance and advantages of conducting compound screening in a multicellular human-based system to address the therapeutic challenges for axon repair and SMA.

## ACKNOWLEDGEMENTS

We thank the SMA Foundation (CJW), Dr. Miriam and Sheldon G Adelson Medical Research Foundation (CJW), Ellen R. and Melvin J. Gordon Center for the Cure and Treatment of Paralysis Award (KH) for financial support. We thank Yiming Zhang for lentivirus preparation from the viral core and Lee Barrett for support in establishing protocols on the Opera Phenix Plus high-content screening platform at the screening core at Boston Children’s Hospital.

## AUTHOR CONTRIBUTIONS

Conceptualization, K.H., and C.J.W.; Methodology, K.H., X.Z., K.C., R.P., and R.P.; Validation, K.H., S.Z., K.Z., A.N., R.M.G., and R.P.; Formal analysis, K.H., X.Z., K.Z., and A.N.; Investigation, K.H., X.Z., S.Z., K.Z., A.N.; Resources, L.R., and C.J.W.; Original draft, K.H., X.Z., and C.J.W.; Supervision, C.J.W.

## SUPPLEMENTARY FIGURE

**Figure S1. NMII inhibition alters muscle morphology without disrupting AChR clustering (A-C)** Quantification of synchronized muscle activity on day 1 and day 3 in myogenic-only cultures (**A**) after DMSO or 25 μM blebbistatin treatment. Sample sizes for **B, C**: day 1 and day 3 myogenic-only, DMSO (n=20), blebbistatin (n=20). Data represents three biological replicates. Stats: Welch’s ANOVA with Dunnett’s T3 post hoc test. Error bar: SEM, *p*-values: indicated in each graph. **(D and E)** Quantification of synchronized activity in myogenic-only cultures treated with DMSO, 1 μM Dantrolene, or 10 μM MPH-220. Number of active myogenic cells (**D**), and synchronized events (**E**) in day 3 cocultures after treatments. Sample sizes for **D, E**: DMSO (n=20), Dantrolene (n=20), MPH-220 (n=20). Data represents three biological replicates. Stats: One-way ANOVA with Tukey’s post hoc test, error bars: SEM. *p*-values: indicated in each graph. **(F)** Cocultures stained for neurofilament (NF), and α-BTX following DMSO or blebbistatin treatment. **(G)** Quantification of α-BTX intensity and myogenic cell thickness normalized to DMSO in cocultures. Sample size: DMSO (n=16), blebbistatin (n=18) for α-BTX intensity, and DMSO (n=15), blebbistatin (n=15) for thickness measurement. Three biological replicates were done, Stats: unpaired *t*-test for α-BTX intensity, and Welch’s *t*-test for thickness measurement, error bars: SEM. *p*-values: indicated in graph.

## METHODS

### Cell culture

#### Human iPSC lines and culture conditions

Human iPSCs (LiPSC-GR1.1) derived from a male newborn donor (Lonza) were used as the healthy control. SMA patient-derived iPSCs were obtained from Lee Rubin’s lab at Harvard (46), including line 15E from a female neonate (SMA type 0, ID#: SMA15), line 1-33L from a 0.7-year-old male (SMA type I, ID#: 1033), and line 1-38D from a 1.6-year-old female (SMA type I, ID#: 1038). G-banded karyotyping was performed on all iPSC lines at WiCell, and all lines displayed a normal karyotype. The cells were maintained in StemFlex medium (ThermoFisher, #A3349401) on Geltrex-coated plates (ThermoFisher, #A1413302). For routine passaging, cells were dissociated using ReLeSR (StemCellTech, #05872).

#### Motor neuron differentiation

hiPSCs were differentiated to motor neurons for 14 days in N2B27 medium (½ DMEM/F12 (ThermoFisher, #11320082) and ½ Neurobasal-A media (Life technologies, #10888-022) supplemented with 1x N2 (Gibco, #17-502-048), 1x B27 (Gibco, #17-504-044), 1x GlutaMAX and 100□μM non-essential amino-acids. During the first 7 days, cells were cultured in N2B27 medium containing 10 μM SB431542 (Cayman Chemical, #13031), 100 nM LDN-193189 (Sigma, #SML0550), 1 μM Retinoic acid (Sigma, #R2625), 1 μM SAG (Tocris, #4366). For the subsequent 7 days, cells were maintained in N2B27 medium with 1 μM Retinoic acid, 1 μM SAG, 4 μM SU5402 (Sigma, #SML0443), and 5 μM DAPT (Tocris, #2634). On day 14, cells are dissociated using accutase and motor neurons are sorted out by magnetic-activated cell sorting (MACS) using anti-human CD56 (NCAM1) antibody (BD Biosciences, #555516) and anti-R-phycoerythrin magnetic particles (BD Biosciences, #557899). Motor neurons were then replated onto plates coated with 25 μg/ml poly-L-ornithine (Sigma, #P4957) and 10□μg/ml laminin (ThermoFisher, #23017015), and maintained in neurobasal-A medium supplemented with 1x N2, 1x B27, 1x GlutaMAX, 100□μM non-essential amino-acids, 35□μg/ml ascorbic acid (Sigma, #A4403), and 10□ng/ml of each of recombinant human BDNF (ThermoFisher, #PHC7074), CNTF (ThermoFisher, #PHC7015), and GDNF (PeproTech, #450-10).

#### Skeletal myogenic cell differentiation

Primary myogenic cell differentiation was performed using healthy donor iPSCs (LiPSC-GR1.1) followed by protocols (47, 48). After dissociation with TrypLE (ThermoFisher, #12563011), 350,000 iPSCs per well were seeded onto Matrigel-coated 6-well plates in mTeSR1 medium (StemCellTech, #85850) with the ROCK inhibitor Y-27632. From day 0 to day 5, cells were maintained in Di medium [DMEM/F12 with Glutamax (ThermoFisher, #10565-018) supplemented with Insulin-Transferrin-Selenium (ITS; Corning, #25-800-CR), Normocin (Invitrogen, #ant-nr-2), and non-essential amino acids (NEAA; ThermoFisher, #11140050)] with 3 μM Chir99021 (Sigma, #SML1046) and 0.5 μM LDN193189 (Tocris, #6053). From day 3 to day 5, 20 ng/ml FGF2 (PeproTech, #AF-100-18B) was added. At day 6, cells were switched to DK medium [DMEM high glucose with Glutamax (ThermoFisher, #10566-016), 15% Knockout serum replacement (ThermoFisher, #10828028), NEAA, 0.1 mM 2-mercaptoethanol, and Normocin] supplemented with 20 ng/ml FGF2, 2 ng/ml IGF-1 (PeproTech, #10011500UG), 10 ng/ml HGF (PeproTech, #100-39H), and 0.5 μM LDN193189. From day 8 to day 20, cultures were maintained in DK medium with 2 ng/ml IGF-1, and from day 12, 10 ng/ml HGF was added.

The cells then transduced with lentivirus expressing MHCK7-GCaMP6f (Addgene, #65042). Secondary differentiation was performed followed by protocol (49). Briefly, 150,000 primarily differentiated myogenic cells per well were seeded in a Matrigel-coated 24-well plate (ibidi, #82426) in skeletal growth media (PromoCell, #C-23060) with Y-27632. The following day, the media was replaced with fresh skeletal growth media, and one day later, cells were fed with DKI media [DMEM high glucose with glutamax, 2% knockout serum replacement, 1x ITS-G (Life Technologies, #41400045), NEAA, normocin] with 1 μM Chir99021, 10 μM SB431542 (Tocris, #1614) and 10 μM Prednisolone (Sigma, #P6004) for 8 days.

#### Neuron-myogenic cell coculture and injury model

To achieve spatial separation of neurons and myogenic cells within the same cultures, 24-well glass-bottom plates were first coated with 25 μg/ml Poly-L-Ornithine in 1.34x borate buffer overnight, following the next day of coating with 10 μg/ml laminin overnight. After drying the plates, silicone inserts (iBidi, #80369) were placed, and the area outside the inserts was coated with Matrigel for 30 min before seeding myoblasts. Inside each insert, 150,000 motor neurons were seeded as spot cultures following established protocols (17), and maintained in motor neuron media supplemented with 10 ng/ml of BDNF, CNTF, and GDNF, 35□μg/ml ascorbic acid. The ROCK inhibitor was removed the next day, and silicone inserts were removed on day 2. Cultures were then maintained in a 1:1 mixture of DKI medium and motor neuron medium supplemented with 10 ng/ml IGF-1, 200 nM ascorbic acid, and 10 ng/ml CNTF and GDNF. On day 6, laser-induced axotomy of axons was performed.

### Compounds

The following compounds were used: 50 μM Tubocurarine chloride pentahydrate (Curare; Sigma, # 93750-250MG), 10 μM MPH-220 (MedChemExpress, #HY-148516), 1 μM dantrolene (Tocris, #0507), 25 μM blebbistatin (Enzo Life Sciences, #BML-EI315-0005).

### Image acquisition and analysis

Time-lapse live-cell imaging and neurite length measurements were performed using the IncuCyte S3 Live-Cell Analysis System (Sartorius). Live-cell GCaMP6f imaging was carried out with a 20× objective and 3×3 binning on the Opera Phenix Plus high-content screening platform (PerkinElmer). For NMJ activity, images were acquired at 10 Hz for 1 min, capturing at least three fields per well and three replicates per condition across biological replicates. Exported images were compiled into combined TIFF files per field using a custom script. Synchronized activity was quantified and visualized as heatmaps or trace maps using modified CaImAn analysis (27). From the heatmap, the maximum number of activated myogenic cells per frame was quantified for each field, and any instance in which more than seven myogenic cells were activated simultaneously was defined as a synchronized event.

### Immunocytochemistry

Cells were fixed with 4% paraformaldehyde (Electron Microscopy Sciences), permeabilized with 0.3% Triton X-100 in PBS, and blocked with 1% BSA in 0.1% Triton X-100 in PBS. Primary antibodies were applied overnight at 4 °C in blocking solution, including anti-ChAT (Millipore, # AB144P), anti-Isl1/2 (DSHB, #39.4D5), anti-neurofilament (Millipore, #AB5539), anti-skeletal myosin-1 (Millipore, # M1570), and α-actinin (Sarcomeric; Sigma, #A7811). The following day, secondary antibodies were incubated overnight at 4 °C, including donkey anti-goat Alexa Fluor 488, (#A11055), donkey anti-mouse Alexa Fluor 568 (ThermoFisher, #A10037), donkey anti-chicken Cy3 (Jackson ImmunoResearch, #703-165-155), and α-bungarotoxin Alexa Fluor 647 (ThermoFisher, #B35450). Nuclei were counterstained with Hoechst 33342 before imaging.

### Statistical measurement

Statistical analyses were performed as follows: for two-group comparisons, unpaired two-tailed Student’s *t*-tests or Welch’s *t*-tests were used. For comparisons among more than two groups, data normality was assessed using the D’Agostino–Pearson omnibus test and variance with Bartlett’s test, followed by one-way ANOVA with Tukey’s post hoc test or Welch’s ANOVA with Dunnett’s T3 post hoc test. For time-course measurements, two-way ANOVA with Tukey’s post hoc test for multiple comparisons was applied. Data are presented as mean□±□SEM. Significance levels are indicated in each graph: \**p*<0.05, \*\**p*<0.01, \*\*\**p*<0.001, \*\*\*\**p*<0.0001. Graphs were plotted using GraphPad Prism.

