## Supplementary figures and images for "A human iPSC-derived motor neuron-myogenic cell coculture platform to evaluate neuromuscular junction innervation after axon injury and in Spinal Muscular Atrophy"

### Supplementary Fig. 1

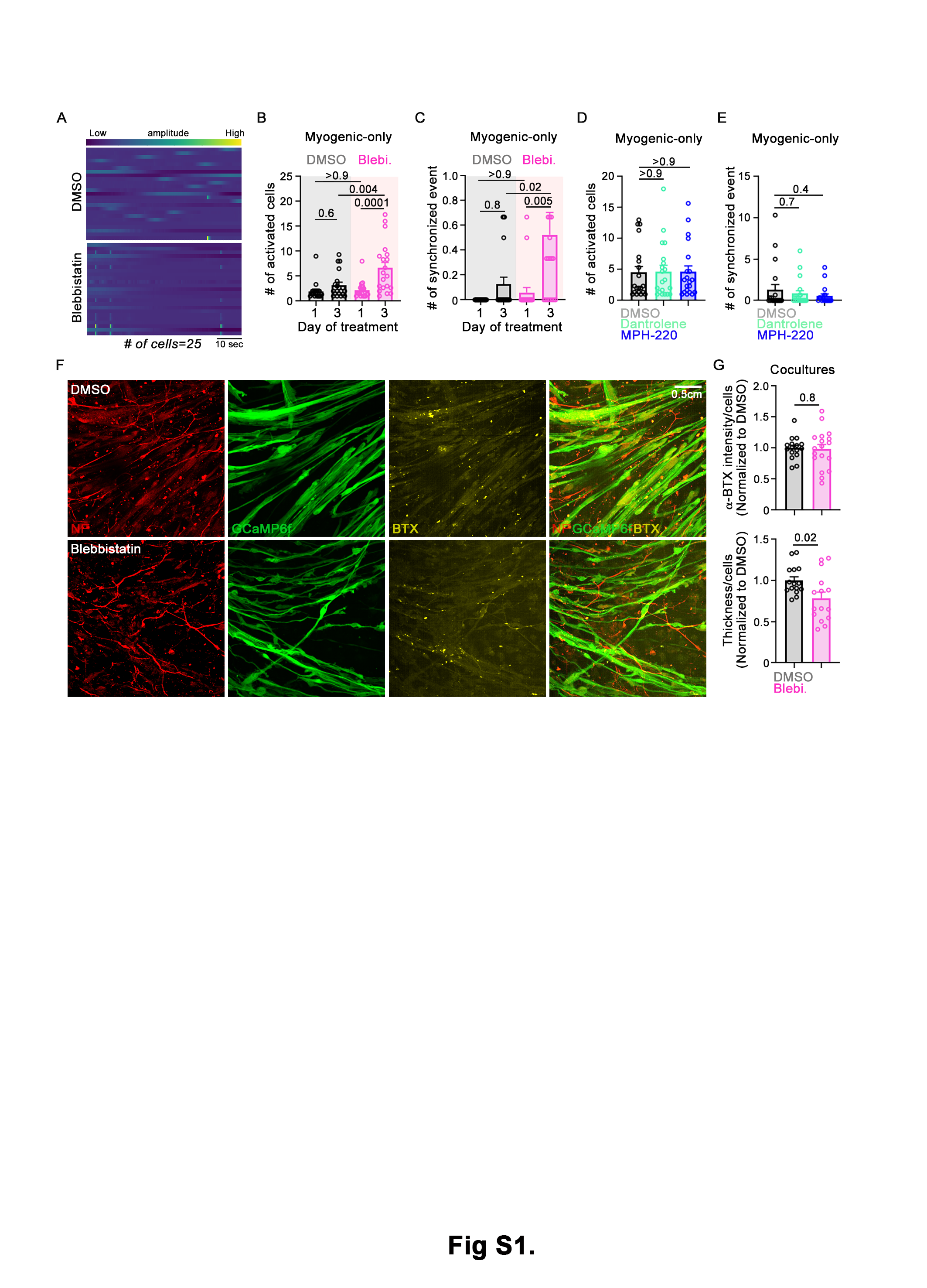
